# Distinct Excitability Properties of Cardiac Calbindin Neurons: Identifying a Unique Neuronal Population

**DOI:** 10.1101/2024.05.31.596786

**Authors:** G. Lizot, J. Bescond, Y. De Koninck, M. Chahine, P. Bois, J-F. Faivre, A. Chatelier

**Affiliations:** PReTI Laboratory, UR 24184, University of Poitiers, Poitiers, France; CERVO Brain Research Centre, Université Laval, Québec, QC, Canada; Department of Medicine, Faculty of Medicine, Université Laval, Quebec City, Quebec, Canada

**Keywords:** intrinsic cardiac neurons, calcium current, Calbindin-D28K, N-type calcium channel, autonomic nervous system

## Abstract

The intrinsic cardiac nervous system is a complex system that plays a critical role in the regulation of cardiac physiological parameters and has been shown to contribute to cardiac arrhythmias. To date, several types of neurons with distinct neurochemical and electrophysiological phenotypes have been identified. However, no study has correlated the neurochemical phenotype to a specific electrophysiological behavior. Calbindin-D28k, a calcium binding protein, is expressed in numerous cardiac neurons. Given that changes in neuronal excitability have been associated with arrhythmia susceptibility and that calbindin expression has been associated with modulations of neuronal excitability, our objective is to assess whether the cardiac calbindin neuronal population has specific properties that could be involved in cardiac modulation and arrhythmias. By using a Cre-Lox mouse model to specifically target calbindin neurons with a fluorescent reporter, we characterized the neurochemical and the electrophysiological phenotype of this cardiac neuronal population. Calbindin neurons exhibit a specific neurochemical profile and a larger soma with shorter neurite length compared to other neurons. This was combined with a distinct electrophysiological signature characterized by a lower excitability with a predominantly phasic profile associated to a lower N-type calcium current density. These properties resemble to the cardiac neuronal remodeling observed in pathologies such as type II diabetes and heart failure. Therefore, we believe that this specific neuronal population deserves investigations in the context of these pathologies.

## 1. Introduction

The nervous control of cardiac functions involves the coordinated action of central and peripheral nervous structures. This coordination results in the modulation of various cardiac parameters including heart rate, contractile force, and electrical conduction velocity. Specifically, these regulatory mechanisms involve a cluster of neurons located within the cardiac tissue forming the intrinsic cardiac nervous system (ICNS). Initially perceived as simple parasympathetic relays, recent studies spanning three decades suggest a more intricate organization of this system. These studies propose that the ICNS is engaged in establishing local and autonomous regulatory loops facilitated by the presence of sensory neurons, interneurons and autonomous neurons ^1^.

Thus, the ICNS emerges as a sophisticated system pivotal in regulating cardiac physiological parameters and has been shown to contribute to cardiac diseases, particularly cardiac arrhythmias. Notably, excessive ICNS activity has been correlated with atrial fibrillation ^2,3^. Ablation of the ganglionated plexus stands out as one therapeutic strategy effectively reducing atrial fibrillation incidence ^4,5^. In addition to atrial electrical disorders, this system has also been implicated in ventricular arrhythmia susceptibility by modulating ventricular refractory periods ^6^. In heart failure, both structural remodeling of the ICNS and electrophysiological alterations of this system driven by changes in ion channel expression levels also increase susceptibility to ventricular arrhythmias ^7,8^. Consequently, the ICNS represents a crucial yet understudied cardiac cell type that intricately regulates both cardiac physiology and physiopathology. To date, various types of neurons exhibiting distinct neurochemical and electrophysiological phenotypes have been recognized. Although most cardiac neurons exhibit a cholinergic phenotype, they can also express markers such as catecholaminergic, nitrergic as well as additional ones like the calcium-binding protein calbindin-D28K, neuropeptide Y (NPY) or the cocaine and amphetamine regulated transcript peptide (CART) ^9–12^.

From an electrophysiological perspective, cardiac neurons can exhibit diverse excitability properties. For example, most cardiac neurons demonstrate a phasic discharge profile, while others are categorized as either accommodating or tonic neurons ^10,13–15^. The rheobase and afterhyperpolarization properties of cardiac neurons are also parameters of discrimination within the cardiac neuron populations ^10^. However, to our knowledge, no study has correlated the neurochemical phenotype with specific electrophysiological properties. Such investigations would elucidate the impact of cardiac neurons on the physiological and pathophysiological functions of the heart. This become increasingly pertinent given that these neurons have been associated in the initiation and perpetuation of cardiac arrhythmias ^3,4,16^.

In a previous study on the mouse heart, we discovered that approximately 46% of cardiac neurons express calbindin-D28k protein ^10^. This neuronal population has also been observed in the rat heart ^11^. Yet, there is currently no information available regarding the specific neurochemical and excitability properties of these neurons. Calbindin-D28K belongs to the family of calcium-binding proteins expressed in various classes of neurons and whose expression serves as a discriminative marker of neuronal populations in both the central and peripheral nervous system ^17–20^. Interestingly, calbindin-D28K protein expression has been correlated with distinct neuronal excitability properties. For instance, calbindin-D28K expression has been linked to alterations in neuronal action potential frequency ^19,21^. Additionally, Calbindin-D28K expression has also been associated with changes in the activity of ionic channels, such as voltage-gated calcium or potassium channels ^21,22^.

Considering the correlation between changes in neuronal excitability and susceptibility to arrhythmias in cardiac pathology ^7,8^ and the association of calbindin expression with modulation of neuronal excitability and ion channels ^19,21,22^, it becomes crucial to investigate whether the cardiac calbindin neuronal population possesses specific properties potentially involved in cardiac modulation. Therefore, employing a Cre-Lox mouse model that enabled precise targeting of calbindin D-28K expressing neurons, our aim is to characterize the neurochemical and electrophysiological properties of this particular cardiac neuronal population.

## 2. Methods

### 2.1 Animals and viral transduction

Experimental procedures were conducted using adult (8-20 weeks) Calb1-IRES2-Cre-D mice (JAX stock #028532). The investigation of calbindin-D28K-expressing cardiac neurons employed a viral transduction strategy utilizing Calb1-IRES2-Cre-D heterozygous mice. Viral transduction was carried out in mice aged between 4 to 6 days. The mice were anesthetized by hypothermia on ice ensuring the mice were adequately sedated. Subsequently, intraperitoneal injection of AAV1 pAAV-FLEX-tdTomato (#28306, Addgene) diluted in 40 µL PBS (6.25.10^12^ VG/mL) was administrated using a 30G needle (Terumo, VWR). The animals were then bred and subsequently used for experimental purposes. All animal procedures performed conform to the guidelines from Directive 2010/63/EU of the European Parliament on the protection of animals used for scientific purposes. The protocol (APAFIS#20949-201906051642993) was ethically approved by the local ethics committee “COMETHEA” (CEEA n°084). For heart excision, mice were intraperitoneally injected with heparin and a combination of Xylasine (10 mg/kg) and Kétamine (80 mg/kg) to anesthetized the mice. Hearts were excised after verification of cessation of pain reflexes.

### 2.2 Immunohistochemistry

Immunohistochemistry experiments were conducted following previously established protocols ^10^. Hearts were promptly excised, rinsed in cold Tyrode solution and fixed in 4% paraformaldehyde for 24h at 4°C. Subsequently, hearts underwent three 3 washes in Phosphate-Buffered Saline (PBS, Corning), followed by cryoprotection in 30% sucrose overnight at 4°C. Prior to sectioning, samples were embedded in OCT tissue compound (Tissue Tek, Sakura Finetek, VWR) and frozen in cold ethanol. Sections 50 µm thick were obtained using a cryostat and mounted onto microscope slides (Adhesion slides, Menzel Gläser, SuperFrost® Plus, Thermo scientific). Sections were washed three times with PBS and permeabilized with 0.5% TritonX-100/1% Bovine Serum Albumin (BSA) in PBS for 2 hours at room temperature. Incubation with primary antibodies (see table 1) was carried out overnight at 4°C. Following three washes with PBS, sections were incubated with secondary antibodies for 3 hours in the dark at room temperature. Nuclei were stained with DAPI (4′,6-diamidino-2-phenylindole, 1:200) and sections were mounted in Mowiol mounting medium. Images were acquired using a confocal laser scanning microscope (FV3000 Olympus, Tokyo, Japan). As a control, images obtained after incubation with the secondary antibodies in the absence of primary antibodies did not elicit any labelling. Quantification was performed by manual cell counting using NIH ImageJ (Bethesda, Maryland, USA). For each marker, quantification was conducted with sections obtained from at least two distinct animals. All micrographs are a projected confocal Z-series from sectioned material.

**Table 1:** Primary and secondary antisera used within this study.

| Primary Antibody | Host Species | Dilution | Catalogue Number | Supplier |
| --- | --- | --- | --- | --- |
| Calbindin D28k | Rabbit | 1:1200 | CB38 | Swant |
| CART (55-102) | Rabbit | 1 :1000 | H-003-62 | Phoenix Pharmaceuticals |
| ChAT | Goat | 1:150 | AB144P | Merck Millipore |
| NPY | Rabbit | 1:500 | Ab10980 | Abcam |
| TH | Chicken | 1:500 | Ab76442 | Abcam |
| tdTomato | Rabbit | 1 :300 | 600-401-379 | Rockland |
| Tuj1 | Rabbit | 1 :200 | Ab52623 | Abcam |
| **Secondary Antibody** |  |  |  |  |
| Anti-chicken (FluoProbe594) | Donkey | 1:300 | FP-SD1110 | Interchim |
| Anti-goat (AF488) | Donkey | 1:300 | A-11055 | ThermoFisher Scientific |
| Anti-rabbit (AF647) | Donkey | 1:300 | A-31573 | ThermoFisher Scientific |

### 2.3 iDISCO

Mouse hearts were stained and cleared using a modified iDISCO+ protocol ^23,24^ as previously described ^10^. Fixed hearts were dehydrated with a graded methanol series (50%, 80% and 100% methanol in PBS, each for 1.5 h at room temperature and bleached overnight with 6% H202 in methanol at 4 °C. Following two washes in 100% methanol, hearts were gradually rehydrated (80%, 50% methanol and PBS, 1 hour and 30 minutes each). Samples were blocked and permeabilized with PBS containing 0.2% gelatin 0.5% triton X-100 and 50mM sodium azide (PBSGT) for 4 days at room temperature. The primary tdTomato antibody (see table 1) was then incubated during two weeks in PBSGT supplemented with 0.1% saponin at 37°C. Subsequently, samples were washed in PBSGT for 1 day followed by a 48 hours incubation with secondary antibodies. After another day of washing, samples were again dehydrated in methanol (20%, 40%, 60, 100% and 100% methanol, 1 hour each) and incubated overnight in a mixture of dichloromethane (DCM) – methanol (2:1) at RT. Samples were incubated in 100% DCM for 30 minutes and finally incubated and stored in DiBenzyl Ether. Imaging was performed by light-sheet microscopy using a UltraMicroscope II (LaVision BioTec, Bielefeld, Germany). Image processing was carried out using Bitplane Imaris 9 software (Oxford instruments company, Abingdon, UK).

### 2.4 Neuronal dissociation

Neuronal dissociation was conducted following the procedure outlined in our prior research ^10^. Hearts were swiftly excised and rinsed in cold HBSS solution. The fat pads located between both atria were dissected, cut into small fragments and dissociated in 2 mL HBSS containing 3 mg/mL collagenase type II (Worthington), 7.5mg/mL dispase II and 0.25 mg/mL DNase I for 30 minutes at 37°C. This was followed by an additional incubation in 2 mL trypsin-EDTA 0.25% supplemented with 0.25 mg/mL DNase I for 35 minutes at 37°C. After two washes in culture media, cells were gently triturated with fire-polished Pasteur pipettes coated with SVF and plated onto laminin-coated 35 mm glass bottom Petri dishes. The cells were maintained in Neurobasal-A medium (ThermoFisher Scientific, Villebon sur Yvette, France) supplemented with 2mM L-glutamine, B27 supplement (ThermoFisher Scientific, Villebon sur Yvette, France), 5% horse serum and 1% penicillin/streptomycin in a humidified chamber at 37°C with 5% CO_2_.

### 2.5 Morphological analysis

The neuronal size was assessed by measuring the soma perimeter and area of both tdTomato positive and negative neurons. These measurements were obtained using NIH ImageJ software on images of neurons utilized in patch-clamp experiments. In the same neurons, the membrane capacitance recorded during patch clamp experiments was used as an indicator of neuronal size. For axonal length measurement, neurons were maintained in the culture medium for 24h to allow axonal growth. Neurons were then fixed with 4% paraformaldehyde in PBS for 10 min, washed (PBS) and permeabilized with 0.1 % TritonX-100 in PBS for 10 min at RT. After washing with PBS, unspecific sites were saturated using PBS containing 1% of BSA and 0.1 % Tween 20 for 30 min and incubated with primary antibody directed against β3-tubulin (Tuj1, see Table 1) diluted in the saturation solution overnight at 4°C. Cells were then washed with PBS and incubated with secondary antibody (see Table 1) for 2 hours in the dark at room temperature. After another PBS wash, cells were mounted in Mowiol mounting medium. Images were captured using a confocal laser scanning microscope (FV3000 Olympus). Neurite length was determined by Tuj1 immunolabelling and quantified using the filament tracer module of Bitplane Imaris 9 software.

### 2.6 Electrophysiology

Electrophysiological properties of isolated cardiac neurons were determined using the whole-cell configuration of the patch clamp technique. Recordings were carried out at room temperature within 24 hours following neurons isolation with Patch electrodes pulled from glass capillaries (PG150T-7.5, Harvard Apparatus, Les Ulis, France) using a vertical micropipette puller (Narishige, Tokyo, Japan). Recordings were made with an Axopatch 200B amplifier (Molecular devices, San Jose, California, USA) with a 5-kHz low-pass filter. Data were sampled at 20 kHz and digitized by a Digidata 1550 B (Molecular devices, San Jose, California, USA). Data acquisition and analysis were performed using pClamp software (v11, Molecular devices, San Jose, California, USA).

For action potential (AP) recordings in current clamp mode, the patch pipettes (≈4 MΩ) were filled with (mM): 130 K-gluconate, 10 KCl, 1 MgCl_2_, 10 HEPES, 1 CaCl_2_, 5 EGTA, 2 Mg-ATP, 10 Na_2_-phosphocreatine and 0.3 Na-GTP, (pH adjusted to 7.2 using KOH). The bath solution contained (mM): 150 NaCl, 5 KCl, 1 MgCl_2_, 1.5 CaCl_2_, 10 glucose, and 10 HEPES, (pH adjusted to 7.4 using NaOH). Liquid junction potential was +16 mV and was not corrected. Input resistance was determined by measuring voltage changes evoked by injection of hyperpolarizing current (from -10 pA to -40 pA; in 10 pA increments). Discharge characteristics were determined by injecting depolarizing current of increasing amplitude (from 20 to 400 pA ; 20 pA increment) during 500 ms. AP properties were determined upon brief injection (2 ms) of suprathreshold current. AP amplitude was measured as the difference between the AP peak amplitude and the resting membrane potential. Spike half-width was calculated as the AP duration measured at 50% of its amplitude.

Voltage clamp experiments were carried out in order to record voltage-gated calcium channel currents. For this purpose, the patch pipettes (3-4 MΩ) were filled with (in mM) 120 CsCl, 10 KCl, 1 MgCl_2_, 40 HEPES, 1 CaCl_2_, 5 BAPTA, 4 Mg-ATP, 0.1 leupeptin, 10 Na_2_-phosphocreatine and 0.3 Tris-GTP, (pH adjusted to 7.3 with CsOH). The bath solution contained (in mM) 140 TEA-Cl, 5 BaCl_2_, 1 MgCl_2_, 2 4-aminopyridine, 10 glucose et 10 HEPES (pH adjusted to 7.4 with TEA-OH). The calculated junction potential of +10,4 mV was corrected before recordings. For Current-voltage relationships, cells were held at -80 mV and a ramp depolarizing protocol was applied from -80 mV to +40 mV during 300 ms. The current voltage relationships were then constructed by reporting the amplitude of the current recorded every 5 mV divided by the cell membrane capacitance measured by the Clampex module of pClamp software. Currents were recorded in the same cell under the perfusion of the bath solution (Control) and with the bath solution containing 1 µM of Omega-conotoxin GVIA (Smartox biotechnology, Saint Egrève, France).

### 2.7 Chemicals

Unless stated, all chemicals were obtained from Sigma-Aldrich (Lyon, France).

### 2.8 Statistical analysis

Statistical analysis was performed using GraphPad Prism (San Diego, California, USA). Data are presented as mean ± SEM. Depending on the data, statistical analysis were performed using either Mann-Whitney, t-test, Fisher exact test or Chi-square tests.

## 3. Results

### 3.1 Neurochemical properties of calbindin neurons

Within the neurochemical diversity of mouse cardiac neurons, our group previously discovered that neurons expressing the calbindin-D28K protein constitute 46% of the total population ^10^. However, until now there has been no information available regarding the neurochemical and electrophysiological properties of this neuronal subpopulation. In order to further advance the investigation of this neuronal population, we opted to employ a Cre-Lox strategy to selectively label these neurons with the tdTomato fluorescent protein. Initially, we confirmed the presence of these neurons in the Calb1-IRES2-Cre-D mice model.

Immunolabelling of both cholinergic and calbindin neurons within cardiac ganglia (Figure 1A) reveals that calbindin neurons constitute a significant portion of ganglionic neurons. Interestingly, calbindin positive projections can form multiple pericellular baskets surrounding neuronal somata (Figure 1A, arrows). These findings align with our previous observations ^10^.

**Figure 1:**
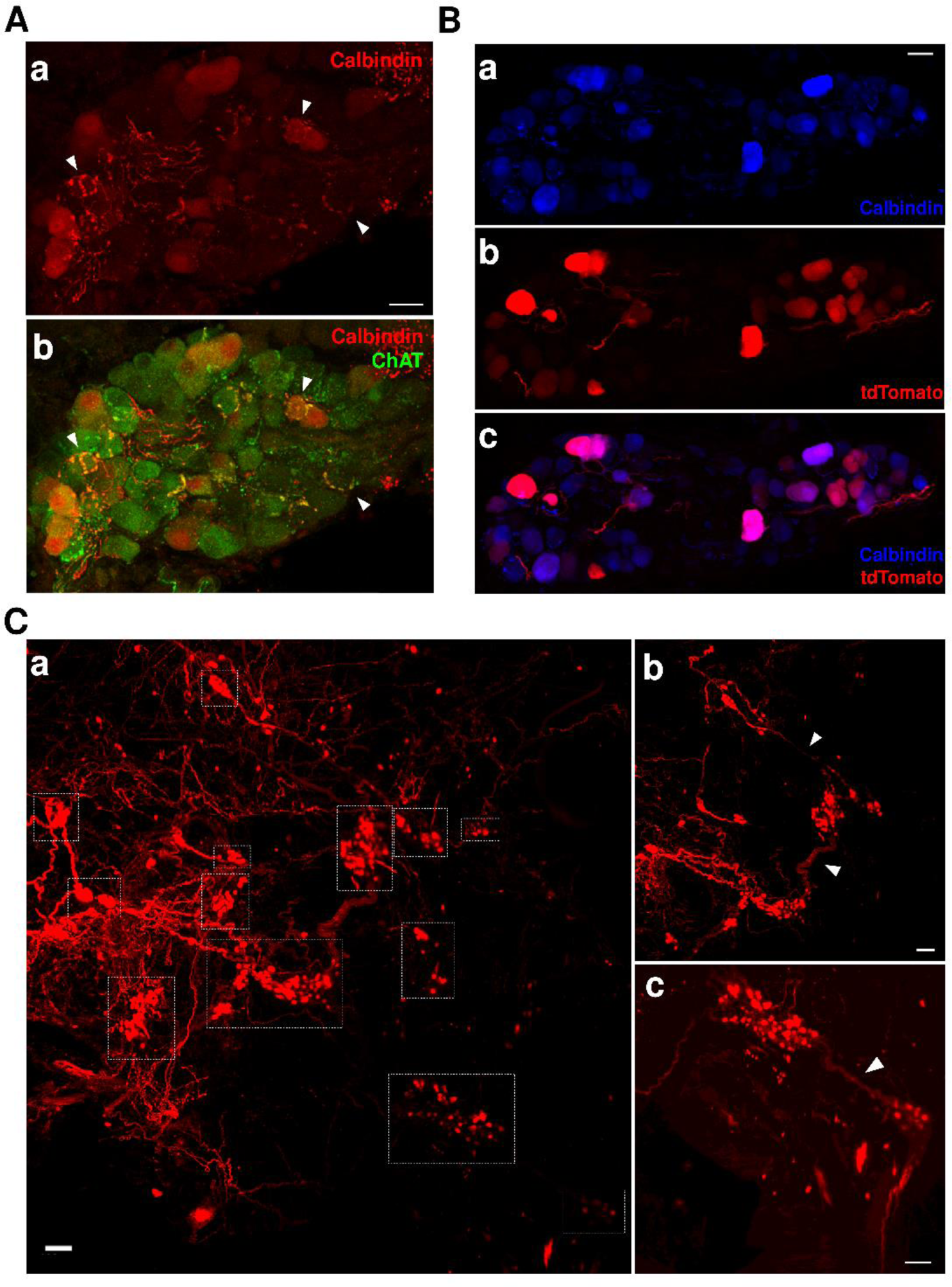
Cre-dependant specific expression tdTomato within calbindin neurons. **A.** Confocal images of sections of a cardiac ganglia immunostained with calbindin (red, **a**). Merged image of the same ganglia showing immunolabelling of both Choline Acetyltransferase (Chat, green) and calbindin (red) neurons **(b)**. Arrows indicate typical calbindin immunoractivity pericellular baskets surrounding neuronal somata (**a, b**). **B.** Confocal images of sections of a cardiac ganglia obtained from AAV1-Flex-tdTomato transduced Calb1-IRES2-Cre-D mice. Calbindin neurons are immunolabelled with calbindin antibodies (blue, **a**) and the tdTomato expression was assessed by red fluorescence (red, **b**). The proportion of calbindin positive neurons (blue) expressing the tdTomato (red) is visible in the merged image (**c**). **C.** Distribution of calbindin neurons within cardiac ganglia assessed by tdTomato immunolabelling in AAV1-Flex-tdTomato transduced Calb1-IRES2-Cre-D mice. **a.** Maximum intensity projection (1926µm) of cleared atria with tdTomato staining (red). The different ganglia are delimited by white rectangles. **b** and **c**. Exemple of ganglia interconnected by calbindin fibers (arrows) identified by tdTomato staining (red). Scale bars are 20 µm (A and B) and 100µm (C).

Therefore, we utilized this mice model to specifically label calbindin neurons, facilitating further investigation into their neurochemical and electrophysiological properties. Through AAV1-Flex-tdTomato viral transduction of Calb1-IRES2-Cre-D mice, we achieved specific expression of tdTomato in calbindin neurons (Figure 1B). This approach resulted in significant expression of the fluorescent reporter in 59 ± 8 % of calbindin-D28K positive neurons (n = 332), leveraging this expression to assess the overall distribution of calbindin neurons within all cardiac ganglia using the iDISCO protocol. Figure 1C shows no preferential localization of these neurons within the cardiac ganglion plexus. Additionally, tdTomato expression is observed in fibers connecting different ganglia suggesting that calbindin neurons could participate in inter-ganglionic communication (Fig1C b,c).

To unravel the neurochemical profile of calbindin neurons, we conducted immunohistochemistry experiments onto tdTomato expressing hearts. This enabled the identification of calbindin neurons by fluorescence, thereby avoiding crossreaction between antibodies. Immunodetection of the cholinergic marker ChAT confirmed that almost all the calbindin neurons are cholinergic (>99%), consistent with previous findings ^10^. Among the phenotypic diversity previously identified in cardiac neurons expressing Tyrosine Hydroxylase (TH), Cocain and Amphetamin Related Transcript peptide (CART) and Neuropeptide Y (NPY) represent significant neuronal populations in mice ^10^. We therefore aimed at evaluating the proportion of expression of these markers within the calbindin neuronal population. 36.5 % of calbindin neurons co-express TH (Figure 2 A. a-c and B.). This proportion is significantly lower (25.9%, p<0.01) in neurons without tdTomato expression referred as “non calbindin neurons” in this study. CART peptide, a neuronal marker expressed in 60% of cardiac neurons ^10^ is expressed in a similar proportion in calbindin neurons (56.9 %) which is comparable to that observed in “non calbindin neurons” (62.4%, p>0.05). However, while the majority of “non calbindin neurons” also express the NPY (70.7%), this proportion was significantly lower (30%, p<0.0001) in calbindin neurons.

**Figure 2:**
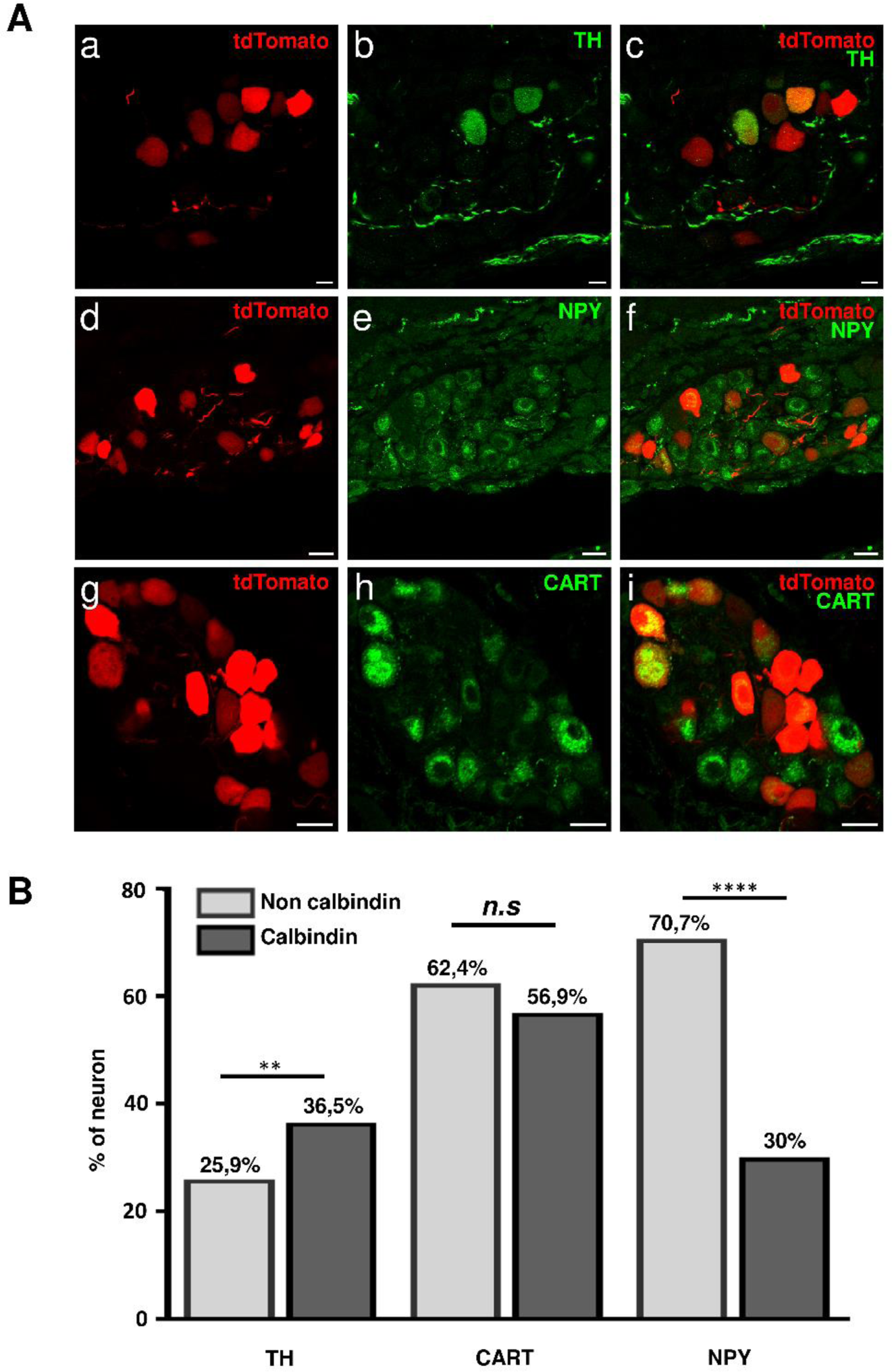
Neurochemical phenotype of calbindin neurons. **A.** Confocal images of cardiac ganglia expressing tdTomato (red) in calbindin neurons and immunostained with either Tyrosin Hydroxylase (TH, green, scale bar 10 µm)) (**a-c**), neuropeptide Y (NPY, green, scale bar 20 µm) (**d-f**) or cocaine and amphetamine related transcript (CART, green, scale bar 20 µm) (**g-f**). **B.** Percentage of neurons expressing TH (n=873), CART (n=950) and NPY (n=371) within the calbindin and non calbindin neuronal population. **(p>0.01) and **** (p<0.0001) indicate significant differences between calbindin and non calbindin neurons (Fisher’s exact test).

**Immunofluorescence experiments reveal that calbindin neurons form a relatively dense network of projections innervating the atrial stage, without being associated with any particular ganglionic zone. This neuronal population exhibits heterogeneity in their neurochemical phenotype, with the co-expression of several markers. However, it is characterized by a slightly higher proportion of TH and a much smaller proportion of NPY expressing neurons.**

### 3.2 Morphological properties of calbindin neurons

The morphological characteristics of calbindin neurons were investigated by examining isolated cells following neuronal dissociation.

Regarding the size of these neurons, calbindin neurons exhibit a larger soma compared to non calbindin neurons (Figure 3, Table 2). This quantification was conducted by measuring the perimeter and area of the soma of isolated neurons (Figure 1A, B). Both parameters are significantly larger in calbindin neurons compared to non calbindin neurons (p<0.01 and p<0.05 respectively for area and perimeter). Hence, calbindin neurons display a soma area (1305.2 ± 120 µm², n=27) that is 41% higher than non calbindin neurons (921.9.1 ± 60.5 µm², n=22). This distinction is further confirmed when comparing the membrane capacitance measured during the whole-cell patch-clamp experiments on these cells. Indeed, cell membrane capacitance is an electrical parameter dependent on the total membrane surface of the cell, thus, reflecting cell volume and size. We observed that calbindin neurons exhibit a significantly higher membrane capacitance (42.3 ± 2.7 pF, p<0.01) increased by 31% compared to non calbindin neurons (32.3 ± 2.3 pF, Figure 3C, Table 2).

**Figure 3:**
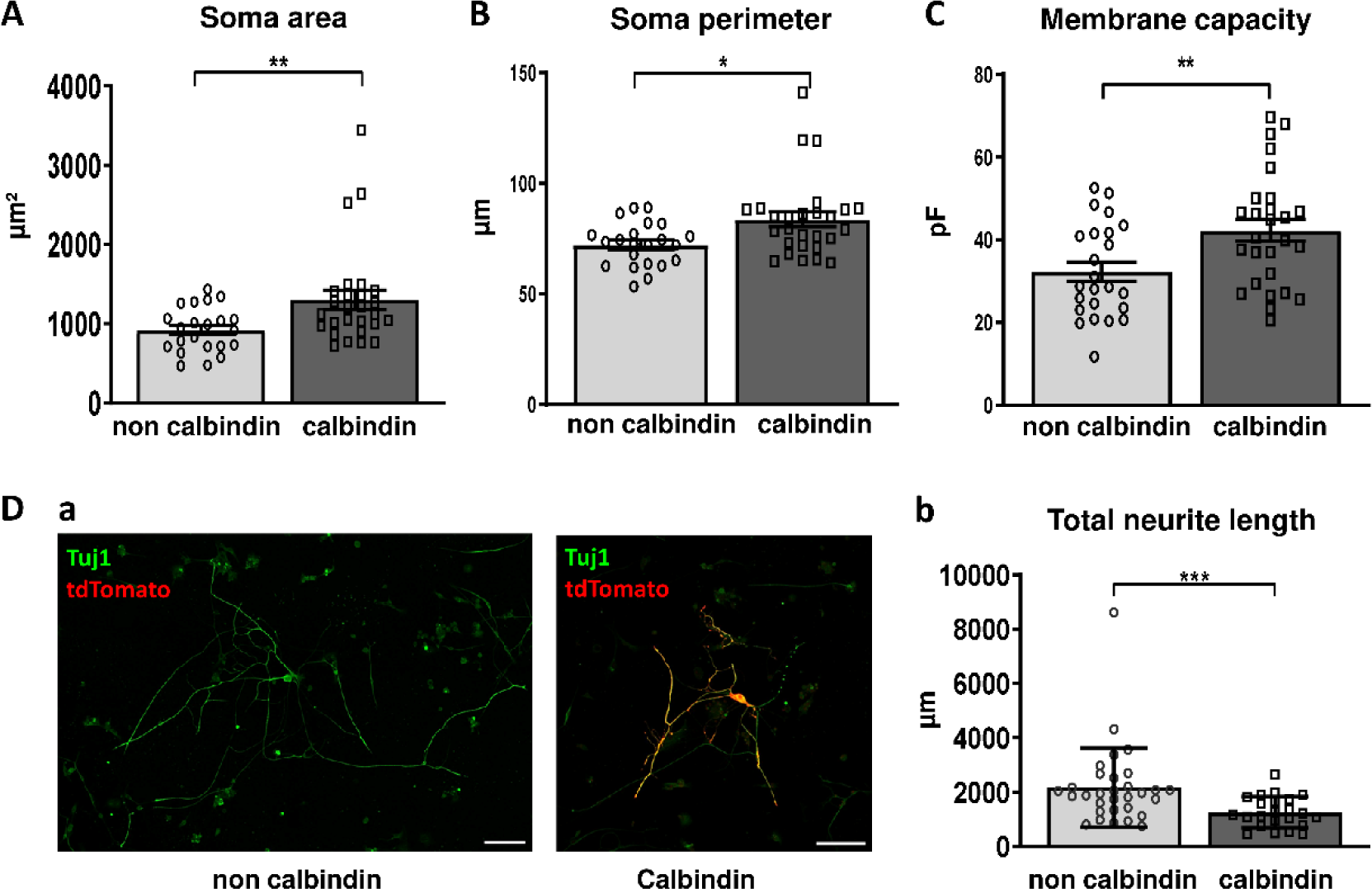
Morphological phenotype of calbindin neurons. Quantification of the neuronal size through measurement of soma area **(A)**, soma perimeter **(B)** and membrane capacitance **(C)** on isolated calbindin and non calbindin neurons. **D.** Neurite lenth assessed on isolated calbindin and non calbindin neurons after 24h of culture. **a.** Example of neurite staining using β-tub III (Tuj1, green) on calbindin (tdtomato, red) and non calbindin neurons. Scale bar: 100 µm. **b.** Total neurite length measured by using the filament tracer module of Imaris software on β-tub III stained calbindin and non calbindin neurons. *** indicates a significant difference (P<0.001, Mann-Whitney; n = 31 (non calbindin) and n = 23 (Calbindin)).

**Table 1.**
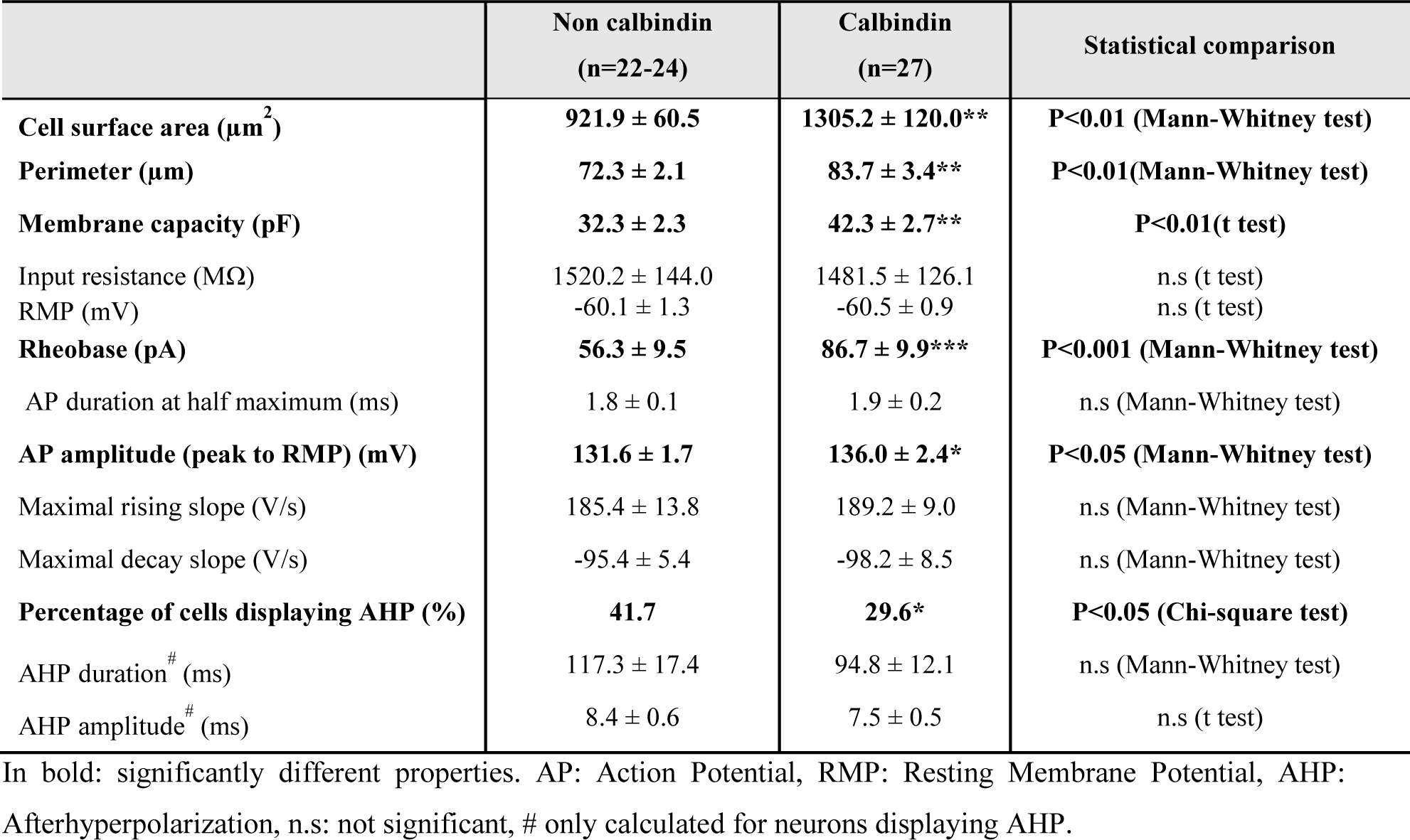
Morphological and electrophysiological properties of non calbindin and calbindin neurons.

We also examined the neurite length of calbindin neurons compared to other neurons. Since most of neurons lose their neurite during cellular dissociation, this parameter was assessed in isolated neurons maintained in culture for 24 hours before fixation and immunolabelling by using β-tubulin III (Figure 3D). These experiments indicate that calbindin neurons exhibit a significantly shorter neurite length (1261 ± 120 µm, n=23, p<0.001) compared to other neurons (2175 ± 262 µm, n=31).

**From a morphological perspective our results suggest that calbindin neurons exhibit a larger soma and a shorter neurite length compared to other neurons.**

### 3.3 Excitability and action potential properties of calbindin neurons

The electrical membrane properties of isolated neurons were assessed using the current clamp mode of the patch clamp technique in whole cell configuration. Calbindin neurons were identified via tdTomato fluorescence and their electrophysiological properties are summarized in Table 2. Regarding passive membrane properties (Figure 4 B, C), the input resistance did not exhibit a significant difference between calbindin (1481 ± 126 MΩ, n=27) and non calbindin neurons (1520 ± 144 MΩ, n=24). Similarly, calbindin neuron displayed a resting membrane potential of -60.5 ± 0.9 mV (n=27), which did not significantly differ from the resting membrane potential of non calbindin neurons (-60.1 ± 1.3 mV, n=24).

**Figure 4:**
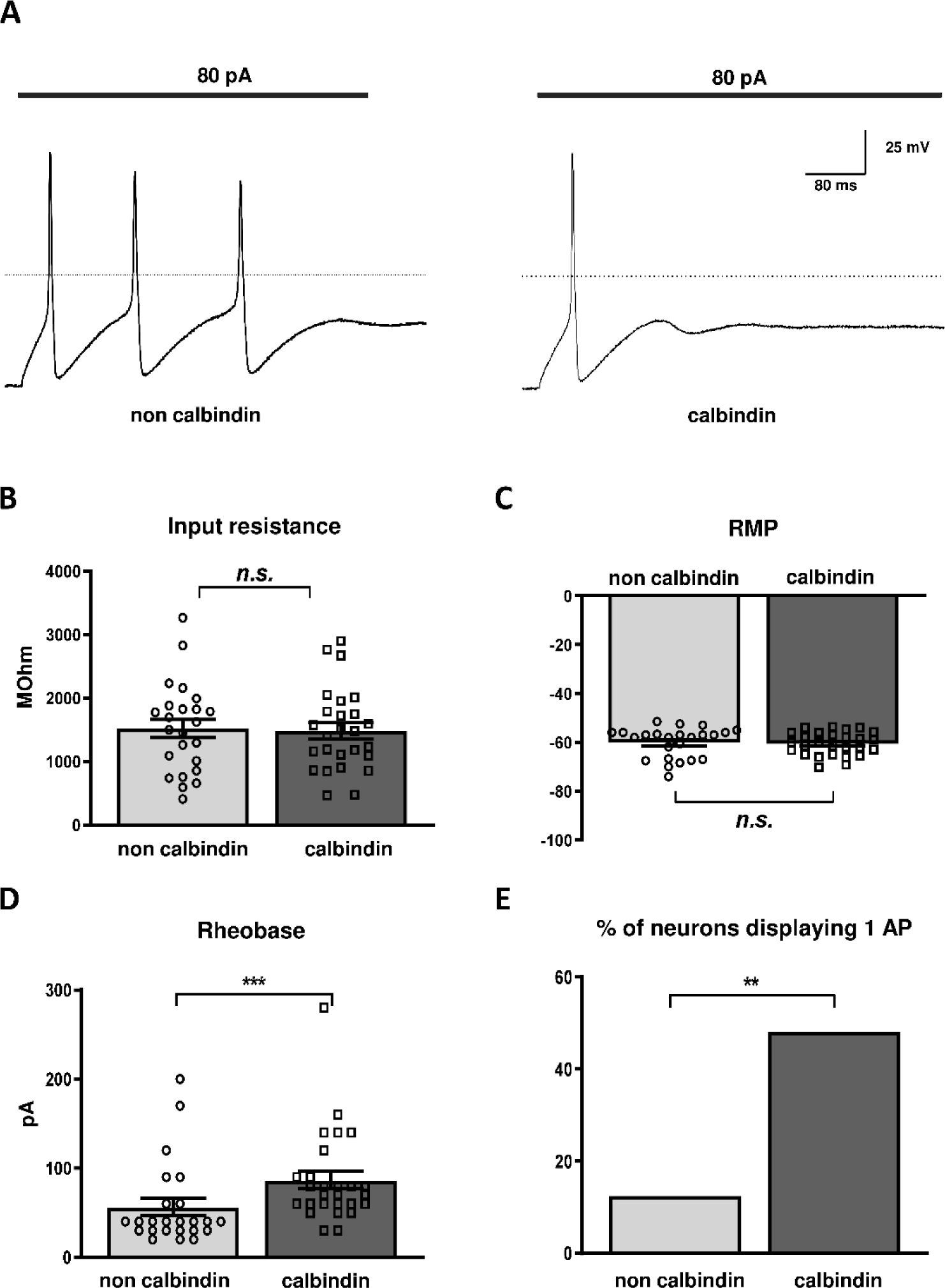
Excitability properties of calbindin neurons. **A.** Representative action potential discharge profile observed in calbindin (right) and non calbindin (left) neurons in response to a 500 ms depolarizing current injection of 80 pA. **B-C.** Comparison of input resistance and Resting Membrane Potential (RMP) recorded in non calbindin and calbindin neurons. **D.** Assessment of the Rheobase of the non calbindin (n=24) and calbindin (n= 27) neurons following a 500 ms depolarizing current injection from 20 to 400 pA (Δ 20 pA). **E.** % of neurons displaying a maximum of only 1 AP recorded in non calbindin (n=24) and calbindin (n=27) neurons with the stimulation protocol used in D. ** (p<0.01) indicates significant differences between non calbindin and calbindin neurons (Fisher’s exact test).

Neuronal excitability (Figure 4 D, E) was evaluated through incremental depolarizing current injection of 500 ms duration. Our findins revealed that calbindin neurons exhibit a significantly higher rheobase (86.7± 9.9 pA, n = 27) compared to non calbindin neurons (56.3 ± 9.5 pA, n=24, p<0.0001). This lower excitability was associated with a reduced discharge capacity of calbindin neurons compared to other neurons. Specifically, regardless of the injected current, 48.15 % of calbindin neurons displayed phasic behaviour, discharging only 1 action potential, in contrast to other neurons where this proportion is significantly smaller (12.5%, p<0.001).

We then sought to determine whether calbindin neurons exhibit specific single AP properties. This was assessed by 2 ms suprathreshold current injections. Similar to the rheobase findings, APs of calbindin neurons are triggered by significantly higher current injection (1142 ± 82 pA, n=27) compared to other neurons (832 ± 74 pA, n=24, p<0.01 Mann-Whitney). Additionally, calbindin neurons are predominantly composed of APs without a hyperpolarization phase (neurons without AHP: 19/27 (calbindin) versus 10/24 (non-calbindin)) (figure 5 A, B, p<0.05). For neurons exhibiting a hyperpolarization phase, the duration and amplitude of this phase were similar for both groups of neurons studied (Table 2). For other parameters of action potential, no significant difference is observed between calbindin and non calbindin neurons in terms of AP amplitude, duration, depolarization and repolarization kinetics (Figure 5 C-F).

**Figure 5.**
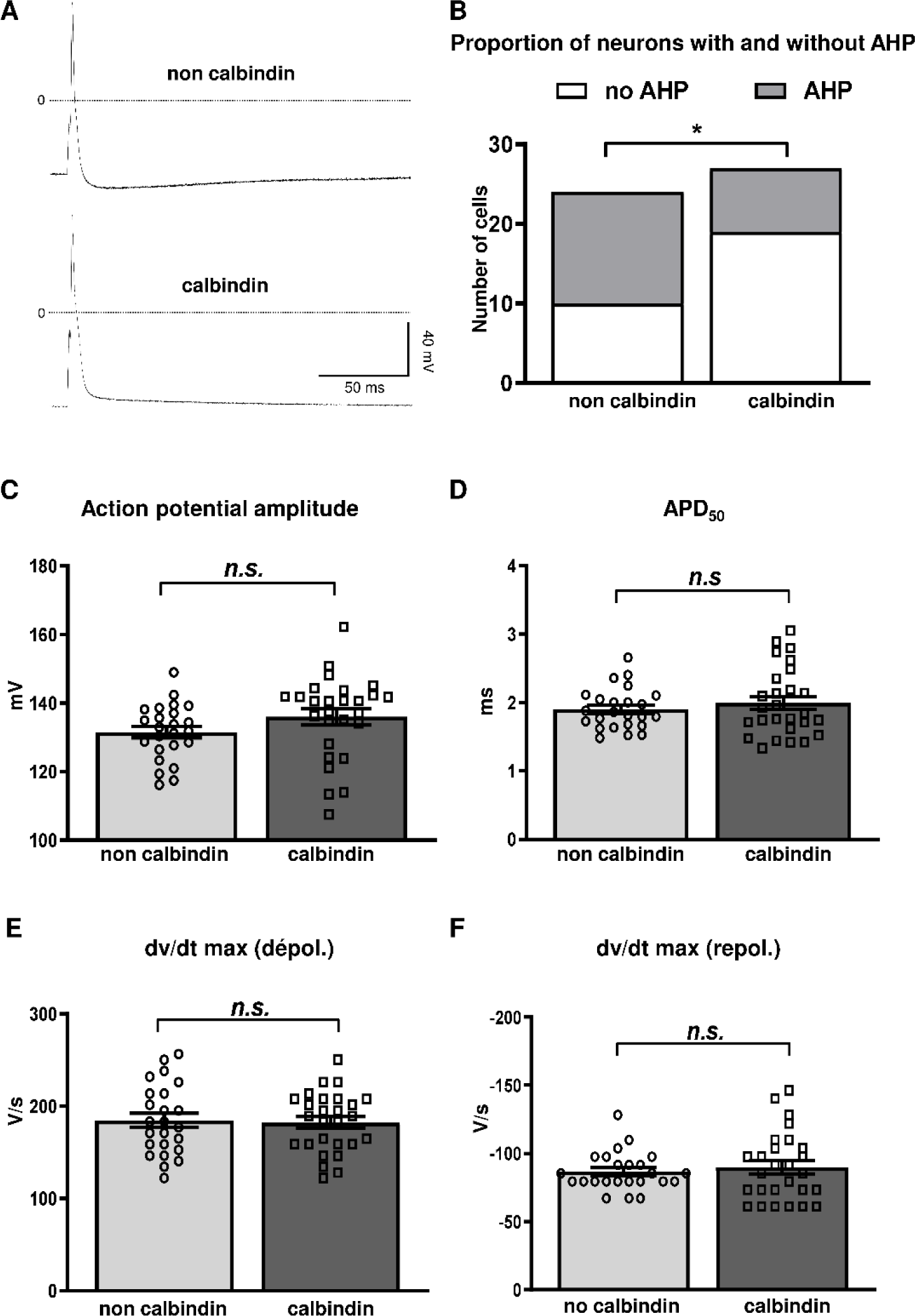
Single action potential properties of calbindin and non calbindin neurons. **A.** representatives action potentials recorded in response to 2 ms depolarizing current injection from 300 to 1500 pA (Δ 40 pA). **B.** Assessment of the proportion of neurons displaying an afterhyperpolarization (AHP) or not (no AHP) in response to the protocol described in A. **C-F.** Action potential amplitude (C), duration (D, APD_50_) and maximum rising (E) and decay (F) slope of neurons. *p<0.05 (Khi-2), *n.s.*: not significant (Mann-Whitney), statistical analysis performed between non calbindin (n=24) and calbindin (n=27) neurons.

**In conclusion, these findings demonstrate that calbindin neurons exhibit a distinct electrophysiological profile marked by a lower excitability, a predominantly phasic firing patterns coupled with a decreased proportion of APs displaying a hyperpolarization phase.**

### 3.4 Calcium current of calbindin neurons

Voltage-gated calcium channels are key players in neuronal excitability ^15,25^. In the context cardiac physiology, inhibition of N-type calcium channels notably leads to an increase in rheobase coupled with a reduction in AP discharge capacity ^25,26^. Since calbindin neurons differ from other cardiac neurons in their low excitability and reduced discharge capacities, we aimed to investigate whether the voltage-dependent calcium currents differed in calbindin neurons. These currents were examined using voltage-clamp techniques with barium as charge carrier.

Under our experimental conditions, depolarization of cardiac neurons triggers an inward current starting at -35 mV and reaching a maximum amplitude around -20 mV in both neuronal populations (Figure 6A). However, the current density is significantly reduced in calbindin-D28K-expressing neurons for potentials between -30 mV and -20 mV (p<0.05). Therefore, we sought to investigate on the same cells the effect of ω-conotoxin, a specific blocker of N-type calcium channels. The perfusion of 1 µM ω-conotoxin results in a marked reduction of current amplitude indicating a significant presence of N-type calcium channels in cardiac neurons (Figure 6B). The ω-conotoxin resistant current of calbindin neurons appears smaller compared to other neurons although this difference was statistically significant only for -20 mV. Hence the ω-conotoxin sensitive current was estimated by the subtraction of the ω-conotoxin resistant current to the global calcium current recorded in the same cell (Figure 6C-D). The N-type current is significantly smaller in calbindin neurons compared to non calbindin neurons with respectively a current density measured at -25 mV of -12.6 ± 3 pA/pF (n=15) and -24.6 ± 3.9 pA/pF (n= 12, p<0.05).

**Figure 6.**
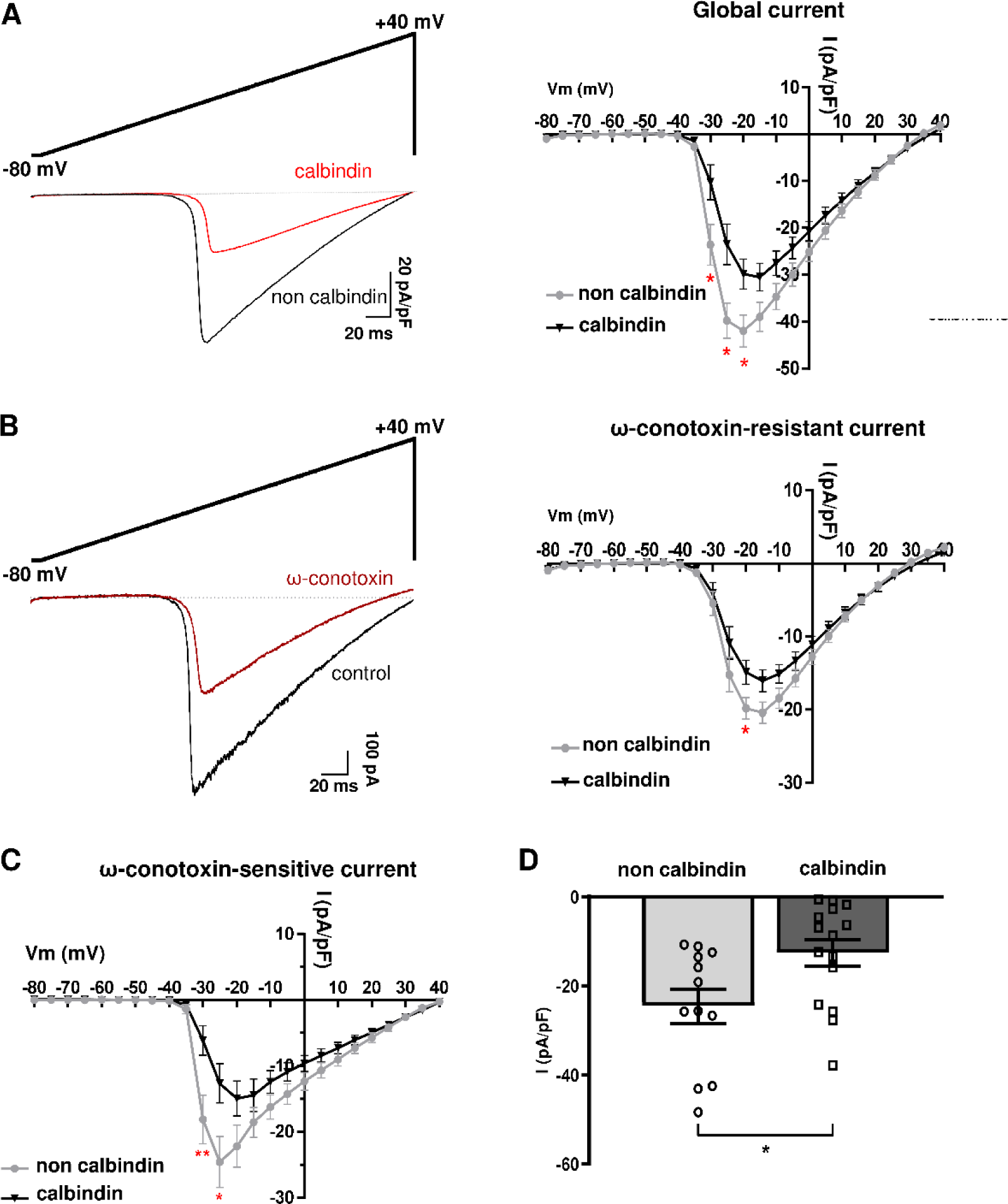
Global and N-type calcium current of calbindin and non calbindin neurons. **A.** Left: Representative traces of global ramp current density obtained in response to a voltage ramp protocol of 300 ms from -80 mV to +40 mV. Right: Current density voltage relationship obtained from the ramp current of non calbindin (black, n= 12) and calbindin (red, n=15) neurons. **B.** Left: Representative trace of the effect of 1µM ω-conotoxin on ramp current obtained in a non calbindin neuron. Right: Current density voltage relationship of the ω-conotoxin resistant ramp current obtained after 1 µM ω-conotoxin perfusion on non calbindin (n= 12) and calbindin (n=15) neurons. **C.** Current density relationship of the ω-conotoxin sensitive current calculated from the difference between global and ω-conotoxin resistant ramp currents in the same neurons. **D.** Bar graph showing the difference of ω-conotoxin sensitive current density recorded at -25 mV in non calbindin (n= 12) and calbindin (n=15) neurons. *p<0.05, **p<0.01 (Student T test).

**Calbindin neurons exhibit a reduced voltage-dependent calcium current density, involving a lower N-type calcium current density. This diminution could account for the less excitable nature observed in these neurons.**

## 4. Discussion

In this study, through the utilization of a Cre/loxP-dependent system for expressing specific fluorescent proteins, we present for the first time an elucidation of the relationship between the neurochemical phenotype, the morphological and electrophysiological properties of cardiac intrinsic neurons. Specifically, our investigation into calbindin-D28k expressing neurons unveils a novel association: That is a minority of these neurons express NPY, featuring enlarged soma with shorter neurite extension, correlated with decreased excitability and a reduced N-type calcium current density. Calbindin-D28K, a calcium-binding protein regulated by vitamin D ^27–29^, has served as a marker to distinguish between neuronal subtypes within both the central and peripheral nervous system ^17–20^. However, the cardiac population of calbindin neurons has been relatively understudied, with only one investigation conducted in rats ^11^ and a prior examination from our team identifying this population in mice ^10^. Our current findings align with those of Richardson *et al*. ^11^ indicating a lack of specific distribution patterns of calbindin neurons across different regions of the ganglionated plexus. Notably, calbindin neurons exhibit both intra- and interganglionic projections, demonstrating the capacity to form pericellular baskets around neuronal somata. These findings suggest that these neurons play a significant role in local circuits. In our investigation, we observed that approximately 70% of non calbindin neurons express the NPY, while this proportion decreased to 30 % within the calbindin population. Since the calbindin population constitutes 45% of the global population in the mouse ^10^, the proportion of neurons co-expressing NPY/calbindin in our study falls within a similar range as observed in rats ^11^. However, in contrast to our results, nearly all neurons, including those within the calbindin population, expressed NPY in rats. This disparity may stem from species differences. In mice, this neuronal population exhibits heterogeneity as evidenced by the fact that the expression of TH, CART and NPY is not strictly confined to either calbindin or non calbindin neurons. Nevertheless, the proportion of TH and NPY is significantly dependent on the presence of calbindin in neurons, suggesting a distinct neurochemical profile for this population.

The morphological profile of calbindin neurons also exhibit notable distinctions compared to other neuronal subtypes. Specifically, they feature a larger soma, as quantified through direct measurements of both area and perimeter, as well as through assessment of membrane capacitance conducted during patch-clamp experiments on the same neurons.

Although this parameter has not been previously explored in calbindin neurons within the heart, a study conducted in the rat stellate ganglion noted larger somas in neurons co-expressing both calbindin-D28K and NPY ^30^. Similarly, in the rat myenteric plexus, calbindin-positive neurons were found to have a larger soma area compared to neurons lacking calbindin-D28K immunoreactivity ^31^. Our study corroborates these findings, demonstrating that the larger size of calbindin neurons is associated with a reduction in neurite length among cultured neurons. This result was derived from a 24 hours culture period subsequent to neuronal dissociation, a necessary step to facilitate neurite outgrowth, as the mechanical dissociation process typically disrupts most neuronal projections. Consequently, the assessment of neurite length under our experimental conditions indicates a reduced capacity for neurite outgrowth among calbindin neurons compared to non calbindin neurons. One plausible explanation is that calbindin-D28k, functioning as a calcium-binding protein, might influence calcium homeostasis, a crucial factor governing axonal growth. ^32,33^.

In addition to their unique neurochemical and morphological characteristics, calbindin neurons, also demonstrate specific electrophysiological properties in our investigation. In a previous study, we delineated four distinct neuronal profiles in mice based on their AP properties ^10^. These profiles included phasic neurons with a higher rheobase in comparison to adapting neurons as well as neurons with or without an AHP. In the present study, the majority of the calbindin neurons fall into the phasic profile, characterized by a single AP with a rheobase significantly higher than that of non calbindin neurons. This was also associated with the absence of AHP in the majority of calbindin neurons. This marks the first instance where the neurochemical profile of a cardiac neuronal population has been linked to its excitability properties. Despite calbindin neurons not constituting a homogenous group, the expression of calbindin is evidently linked to reduced excitability. Additionally, the N-type calcium current density is also significantly lower in this population, potentially accounting for this distinction. Previous studies have associated N-type calcium current with the excitability of cardiac neurons, albeit without discerning between different populations ^25,26,34^.

Exploring the correlation between calbindin expression and neuronal excitability, as well as ion channel expression, has been a focus in various cellular models. For example, it has been shown that dentate granule cell excitability is increased in a calbindin-D28K knockout mouse model ^21^, an effect attributed to a reduction in voltage-gated potassium currents. Similarly, calbindin-D28K expression in a rat beta-cell line results in a reduction of voltage-gated calcium currents ^22^. These findings are in alignment with our observations and suggests that the disparities observed in cardiac neurons may directly stem from the presence of calbindin-D28K protein. This underscores the potential role of calbindin in modulating ion channel activity and, consequently, neuronal excitability.

The findings regarding the properties of cardiac calbindin neurons uncovered in our study hold particular interest with respect to their potential involvement in cardiac physiopathology. Notably, these neurons exhibit an enlarged soma size coupled with a reduced excitability with a lower N-type calcium channel expression. Such characteristics bear resemblance to those observed in cardiac neuronal remodeling associated with conditions such as heart failure and type II diabetes. Indeed, previous studies have noted comparable features in cardiac neurons from patients with heart failure, wherein a hypertrophied cell body was observed in contrast to neurons from healthy hearts ^7^. Corresponding observations have also been documented in this study in animal models of heart failure. Because of the expanded membrane surface area observed, Singh’s group ^7^ suggested that this hypertrophy might contribute to a decrease in neuronal excitability and potentially elucidating the parasympathetic dysfunction linked with heart failure. Indeed, in a rat model of chronic heart failure, cardiac neurons displayed reduced excitability, attributed to a decrease in the N-type calcium channel expression ^35^. This electrical remodeling of intracardiac neurons has been linked to increased susceptibility to ventricular arrhythmias ^8,25^. Such insights underscore the intricate interplay between neuronal remodeling and cardiac dysfunction in pathological conditions like heart failure.

Similarly, in a rat model of type II diabetes, arrhythmia susceptibility was correlated with reduced N-type calcium currents, resulting in decreased neuronal excitability ^26,34^. This remodeling mirrors the characteristics observed in cardiac calbindin neurons, prompting an important inquiry into whether changes in calbindin expression could underlie these pathologies. Furthermore, calbindin D28K expression is known to be dependent on vitamin D3 ^27–29^. Since vitamin D deficiency has been associated with type 2 diabetes, heart failure, and atrial fibrillation ^36^, investigating calbindin expression in cardiac neurons within this context holds considerable interest. Such exploration could potentially uncover novel insights into the role of calbindin in the pathophysiology of these conditions and its implications for neuronal function in the heart

## Limitation of the study

In our experimental setup, AAV1-Flex-tdTomato viral transduction in Calb1-IRES2-Cre-D mice, leads to an expression of the tdTomato in approximatively 60% of calbindin positive neuron. This implies that, while all tdTomato neurons are calbindin positive, there remains a proportion of neurons lacking tdTomato expression (estimated at around 25%) that are still calbindin-positive. Consequently, the discrepancies observed in our study are inevitably underestimated due to this incomplete labeling of calbindin-positive neurons.

## Conclusion

Our current investigation characterizes for the first time the electrophysiological properties of a particular cardiac neuronal population delineated by its neurochemical phenotype. Calbindin neurons exhibit distinct morphological and excitability traits consistent with cardiac neuronal remodeling observed in conditions like type II diabetes and heart failure. Given these parallels, it is imperative to explore further this specific neuronal subset within the context of such pathologies. By doing so, we may gain deeper insights into the complex mechanisms underlying cardiac dysfunction and potentially uncover novel therapeutic avenues.

## Funding

This work was supported by the Fondation pour la Recherche Médicale [FRM DPC20171138946, FDT202106012947] and the UP-SQUARED program of National Research Agency [ANR-21-EXES-0013].

## Acknowledgments

This work has benefited from the facilities of ImageUP and PREBIOS platforms (University of Poitiers) and the technical assistance of Anne Cantereau, Christophe Magaud and Isabelle Fixe.

## Conflict of interest

The authors declare that they have no conflict of interest.

## Data availability

The data underlying this article will be shared on reasonable request to the corresponding author.

## Notes

### Competing Interest Statement

The authors have declared no competing interest.

### Summary of Updates

Figure 3, 4, 5, 6 revised

## References

1. Armour JA. The little brain on the heart. Cleve Clin J Med 2007;74 Suppl 1:S48–51.

2. Ashton JL, Prince B, Sands G, Argent L, Anderson M, Smith JEG, Tedoldi A, Ahmad A, Baddeley D, Pereira AG, Lever N, Ramanathan T, Smaill BH, Montgomery JM. Electrophysiology and 3D-imaging reveal properties of human intracardiac neurons and increased excitability with atrial fibrillation. J Physiol 2024.

3. Choi E-K, Shen MJ, Han S, Kim D, Hwang S, Sayfo S, Piccirillo G, Frick K, Fishbein MC, Hwang C, Lin S-F, Chen P-S. Intrinsic cardiac nerve activity and paroxysmal atrial tachyarrhythmia in ambulatory dogs. Circulation 2010;121:2615–2623.

4. Avazzadeh S, McBride S, O’Brien B, Coffey K, Elahi A, O’Halloran M, Soo A, Quinlan LR. Ganglionated Plexi Ablation for the Treatment of Atrial Fibrillation. J Clin Med 2020;9:E3081.

5. Lemola K, Chartier D, Yeh Y-H, Dubuc M, Cartier R, Armour A, Ting M, Sakabe M, Shiroshita-Takeshita A, Comtois P, Nattel S. Pulmonary vein region ablation in experimental vagal atrial fibrillation: role of pulmonary veins versus autonomic ganglia. Circulation 2008;117:470–477.

6. Jungen C, Scherschel K, Eickholt C, Kuklik P, Klatt N, Bork N, Salzbrunn T, Alken F, Angendohr S, Klene C, Mester J, Klöcker N, Veldkamp MW, Schumacher U, Willems S, Nikolaev VO, Meyer C. Disruption of cardiac cholinergic neurons enhances susceptibility to ventricular arrhythmias. Nat Commun 2017;8:14155.

7. Singh S, Sayers S, Walter JS, Thomas D, Dieter RS, Nee LM, Wurster RD. Hypertrophy of neurons within cardiac ganglia in human, canine, and rat heart failure: the potential role of nerve growth factor. J Am Heart Assoc 2013;2:e000210.

8. Zhang D, Tu H, Wang C, Cao L, Muelleman RL, Wadman MC, Li Y-L. Correlation of Ventricular Arrhythmogenesis with Neuronal Remodeling of Cardiac Postganglionic Parasympathetic Neurons in the Late Stage of Heart Failure after Myocardial Infarction. Front Neurosci 2017;11:252.

9. Horackova M, Armour JA, Byczko Z. Distribution of intrinsic cardiac neurons in whole-mount guinea pig atria identified by multiple neurochemical coding. A confocal microscope study. Cell Tissue Res 1999;297:409–421.

10. Lizot G, Pasqualin C, Tissot A, Pagès S, Faivre J-F, Chatelier A. Molecular and functional characterization of the mouse intrinsic cardiac nervous system. Heart Rhythm 2022:S1547-5271(22)01898–7.

11. Richardson RJ, Grkovic I, Anderson CR. Immunohistochemical analysis of intracardiac ganglia of the rat heart. Cell Tissue Res 2003;314:337–350.

12. Richardson RJ, Grkovic I, Anderson CR. Cocaine- and amphetamine-related transcript peptide and somatostatin in rat intracardiac ganglia. Cell Tissue Res 2006;324:17–24.

13. Hoard JL, Hoover DB, Wondergem R. Phenotypic properties of adult mouse intrinsic cardiac neurons maintained in culture. Am J Physiol Cell Physiol 2007;293:C1875–1883.

14. Xi X, Randall WC, Wurster RD. Electrophysiological properties of canine cardiac ganglion cell types. J Auton Nerv Syst 1994;47:69–74.

15. Xu ZJ, Adams DJ. Resting membrane potential and potassium currents in cultured parasympathetic neurones from rat intracardiac ganglia. J Physiol 1992;456:405–424.

16. Gibbons DD, Southerland EM, Hoover DB, Beaumont E, Armour JA, Ardell JL. Neuromodulation targets intrinsic cardiac neurons to attenuate neuronally mediated atrial arrhythmias. Am J Physiol Regul Integr Comp Physiol 2012;302:R357–364.

17. Ding L, Balsamo G, Chen H, Blanco-Hernandez E, Zouridis IS, Naumann R, Preston-Ferrer P, Burgalossi A. Juxtacellular opto-tagging of hippocampal CA1 neurons in freely moving mice. eLife 2022;11:e71720.

18. Furness JB, Trussell DC, Pompolo S, Bornstein JC, Smith TK. Calbindin neurons of the guinea-pig small intestine: quantitative analysis of their numbers and projections. Cell Tissue Res 1990;260:261–272.

19. Huang S, Egan JM, Fry WM. Electrophysiological properties of rat subfornical organ neurons expressing calbindin D28K. Neuroscience 2019;404:459–469.

20. Smolilo DJ, Costa M, Hibberd TJ, Brookes SJH, Wattchow DA, Spencer NJ. Distribution, projections, and association with calbindin baskets of motor neurons, interneurons, and sensory neurons in guinea-pig distal colon. J Comp Neurol 2019;527:1140–1158.

21. Kim K-R, Jeong H-J, Kim Y, Lee SY, Kim Y, Kim H-J, Lee S-H, Cho H, Kang J-S, Ho W-K. Calbindin regulates Kv4.1 trafficking and excitability in dentate granule cells via CaMKII-dependent phosphorylation. Exp Mol Med 2021;53:1134–1147.

22. Lee D, Obukhov AG, Shen Q, Liu Y, Dhawan P, Nowycky MC, Christakos S. Calbindin-D28k decreases L-type calcium channel activity and modulates intracellular calcium homeostasis in response to K+ depolarization in a rat beta cell line RINr1046-38. Cell Calcium 2006;39:475–485.

23. Belle M, Godefroy D, Couly G, Malone SA, Collier F, Giacobini P, Chédotal A. Tridimensional Visualization and Analysis of Early Human Development. Cell 2017;169:161–173.e12.

24. Renier N, Adams EL, Kirst C, Wu Z, Azevedo R, Kohl J, Autry AE, Kadiri L, Umadevi Venkataraju K, Zhou Y, Wang VX, Tang CY, Olsen O, Dulac C, Osten P, Tessier-Lavigne M. Mapping of Brain Activity by Automated Volume Analysis of Immediate Early Genes. Cell 2016;165:1789–1802.

25. Zhang D, Tu H, Cao L, Zheng H, Muelleman RL, Wadman MC, Li Y-L. Reduced N-Type Ca2+ Channels in Atrioventricular Ganglion Neurons Are Involved in Ventricular Arrhythmogenesis. J Am Heart Assoc 2018;7:e007457.

26. Liu J, Tu H, Zheng H, Zhang L, Tran TP, Muelleman RL, Li Y-L. Alterations of calcium channels and cell excitability in intracardiac ganglion neurons from type 2 diabetic rats. Am J Physiol Cell Physiol 2012;302:C1119–1127.

27. Acharya M, Singh N, Gupta G, Tambuwala MM, Aljabali AAA, Chellappan DK, Dua K, Goyal R. Vitamin D, Calbindin, and calcium signaling: Unraveling the Alzheimer’s connection. Cellular Signalling 2024;116:111043.

28. Alexianu ME, Robbins E, Carswell S, Appel SH. 1Alpha, 25 dihydroxyvitamin D3-dependent up-regulation of calcium-binding proteins in motoneuron cells. J Neurosci Res 1998;51:58–66.

29. Brimblecombe KR, Vietti-Michelina S, Platt NJ, Kastli R, Hnieno A, Gracie CJ, Cragg SJ. Calbindin-D28K Limits Dopamine Release in Ventral but Not Dorsal Striatum by Regulating Ca ^2+^ Availability and Dopamine Transporter Function. ACS Chem Neurosci 2019;10:3419–3426.

30. Richardson RJ, Grkovic I, Allen AM, Anderson CR. Separate neurochemical classes of sympathetic postganglionic neurons project to the left ventricle of the rat heart. Cell Tissue Res 2006;324:9–16.

31. Masliukov PM, Moiseev K, Budnik AF, Nozdrachev AD, Timmermans J-P. Development of Calbindin- and Calretinin-Immunopositive Neurons in the Enteric Ganglia of Rats. Cell Mol Neurobiol 2017;37:1257–1267.

32. Tang F, Dent EW, Kalil K. Spontaneous Calcium Transients in Developing Cortical Neurons Regulate Axon Outgrowth. J Neurosci 2003;23:927–936.

33. Yang Y, Yu Z, Geng J, Liu M, Liu N, Li P, Hong W, Yue S, Jiang H, Ge H, Qian F, Xiong W, Wang P, Song S, Li X, Fan Y, Liu X. Cytosolic peptides encoding CaV1 C-termini downregulate the calcium channel activity-neuritogenesis coupling. Commun Biol 2022;5:484.

34. Zhang D, Tu H, Hu W, Duan B, Zimmerman MC, Li Y-L. Hydrogen Peroxide Scavenging Restores N-Type Calcium Channels in Cardiac Vagal Postganglionic Neurons and Mitigates Myocardial Infarction-Evoked Ventricular Arrhythmias in Type 2 Diabetes Mellitus. Front Cardiovasc Med 2022;9:871852.

35. Tu H, Liu J, Zhang D, Zheng H, Patel KP, Cornish KG, Wang W-Z, Muelleman RL, Li Y-L. Heart failure-induced changes of voltage-gated Ca2+ channels and cell excitability in rat cardiac postganglionic neurons. Am J Physiol Cell Physiol 2014;306:C132–142.

36. Cosentino N, Campodonico J, Milazzo V, De Metrio M, Brambilla M, Camera M, Marenzi G. Vitamin D and Cardiovascular Disease: Current Evidence and Future Perspectives. Nutrients 2021;13:3603.

